# Can you *feel* what I am saying? Speech-based vibrotactile stimulation enhances the cortical tracking of attended speech in a multi-talker background

**DOI:** 10.1101/2025.10.31.685484

**Authors:** I. Sabina Răutu, Mathieu Bourguignon, Vincent Wens, Veikko Jousmäki, Julie Bertels, Xavier De Tiège

## Abstract

In environments with multiple talkers, humans can ‘tune in’ to a speaker of interest while ignoring competing voices. In such conditions, however, auditory cortices track the attended speech envelope rhythms (cortical tracking of speech, CTS) less accurately than in quiet, hindering intelligibility. Visual speech cues (e.g., lip movements) can enhance this CTS, but it remains unclear whether other non-auditory sensory cues, such as tactile input, provide comparable benefits through similar neural mechanisms. Here, using magnetoencephalography, we quantified syllabic (4-8 Hz) and phrasal (<1 Hz) CTS while participants attended to connected speech alone, together with synchronous or asynchronous speech-based vibrations, or with the corresponding speaker video, in both quiet and a multi-talker background noise. We hypothesized that, in noise, speech-based vibrotactile stimulation improves comprehension by enhancing CTS and modulating auditory-seeded functional brain connectivity with extra-auditory neocortical brain areas. Results revealed that synchronous vibrotactile stimulation improved comprehension in multi-talker noise and increased syllabic CTS at right auditory cortex, with this CTS increase magnitude correlating with comprehension performance. Audio-tactile CTS enhancement was accompanied by stronger beta-band auditory cortex connectivity with ipsilateral angular and ventral inferior temporal gyri, alongside reduced alpha-band coupling with the precuneus. These findings suggest that vibrotactile input can support speech-in-noise processing by impacting both local auditory cortical activity and auditory-seeded long-range functional connectivity.

## Introduction

Noisy social environments with simultaneous talkers, such as crowded classrooms or busy cafés, create challenging acoustic conditions for speech comprehension. When faced with this scenario, listeners often engage demanding high-level perceptual and cognitive processes^1–3^ to effectively attend to the speaker of interest and tune out competing talkers. Visual speech cues (e.g., articulatory facial gestures) can reinforce the perceptual segregation of the attended speaker from such a multi-talker background^4,5^, improving intelligibility. Intriguingly, tactile stimulation conveying speech-derived cues can also robustly bolster speech recognition in such environments, even in untrained, normal-hearing listeners^6,7^. Still, the precise neural substrates underlying this audio-tactile enhancement remain to be fully elucidated.

When exposed to connected speech, auditory cortical rhythms synchronize with the slow (<8 Hz) amplitude fluctuations of the speech temporal envelope^8–11^. This cortical tracking of speech (CTS), thought to underlie speech recognition by parsing speech input into its hierarchical linguistic constituents, such as syllables/words (∼4–8 Hz) and phrases/sentences (<1 Hz)^12,13^, is critical for intelligibility^13,14^. In multi-talker settings, CTS selectively aligns with the attended speaker’s speech envelope rather than those of background speakers^2,4,15^, but this selective tracking is less accurate than in quiet, even when the attended speech is louder than the background noise^2,16,17^. Access to the attended speaker’s visual speech cues can effectively enhance CTS^4,16–18^, providing a mechanistic foundation for the behavioral benefits of visual speech in multi-talker backgrounds. Evidence of tactile speech-in-noise enhancement^6,7,19^ suggests that such cross-modal CTS support may generalize to the somatosensory modality. If so, akin to visual cues, speech-derived tactile inputs could support speech recognition in multi-talker noise by honing the CTS of the attended speaker, a supposition that has not yet been investigated.

At the neural level, tactile input can shape auditory processes through a direct modulation of activity in auditory cortices^20–25^, including cortical speech responses^26^. Vibrotactile stimulation transmitting speech-related cues has even been shown to enhance CTS of degraded speech^27^ and of speech embedded in spectrally-matched noise^28^. Still, it remains uncertain whether comparable audio-tactile benefits would emerge in the more ecologically valid context of actively segregating a target speaker’s stream from competing ones. Moreover, the limited spatial resolution conferred by the electroencephalographic (EEG) recordings of these prior studies precluded a precise localization of these effects to the auditory cortex. Rather than more accurate speech representations in the auditory cortex, the observed audio-tactile CTS enhancements may partially reflect the processing of the vibrotactile input in extra-auditory regions (i.e., somatosensory and multisensory). In addition, cross-modal CTS improvements localized to the auditory cortex could be driven by top-down influences from high-order, associative cortical areas through functional connectivity changes, as demonstrated for audio-visual speech perception^29^. The involvement of a more widespread network during audio-tactile conditions is supported by evidence of functional connectivity changes extending beyond auditory cortex following audio-tactile speech training^30^. Nonetheless, the distinct contributions of auditory and extra-auditory activity to audio-tactile CTS enhancement are still largely unknown.

In the present study, we investigated whether CTS enhancement is a core mechanism underlying audio-tactile speech benefits in multi-talker settings. We analyzed magnetoencephalographic (MEG) recordings acquired from thirty normal-hearing adults while they watched four videos containing naturalistic spoken narratives, lasting approximately 6 min each (Figure 1). Speech was delivered either in quiet or mixed with a multi-talker background of equal intensity (i.e., signal to noise ratio (SNR): 0 dB) and presented under three sensory conditions: unimodal (auditory-only, A-only), paired with a unimanual (left hand) speech-derived vibrotactile stimulation (audio-tactile, AT) previously shown to improve speech recognition in multi-talker noise^6^, or accompanied by a video of the speaker’s talking face (audio-visual, AV). This design allowed us to additionally investigate if AT benefits rely on similar CTS-based mechanisms as in AV conditions. To determine whether temporal alignment is obligatory for audio-tactile CTS enhancement, we also manipulated the vibrations’ temporal congruency, contrasting multi-talker conditions in which they were aligned (congruent AT condition; ATc) or misaligned (incongruent AT condition; ATi) with the attended speech envelope. CTS was quantified by computing the coherence between source-localized signals in bilateral auditory cortex and the attended speech envelope at syllabic (4–8 Hz) and phrasal (0.5 Hz) timescales. Critically, we established a direct link between neural-level changes in CTS and behavior by employing condition-specific assessments of comprehension using content questions. Finally, we examined functional brain connectivity changes seeded from auditory cortices involved in CTS.

**Figure 1.**
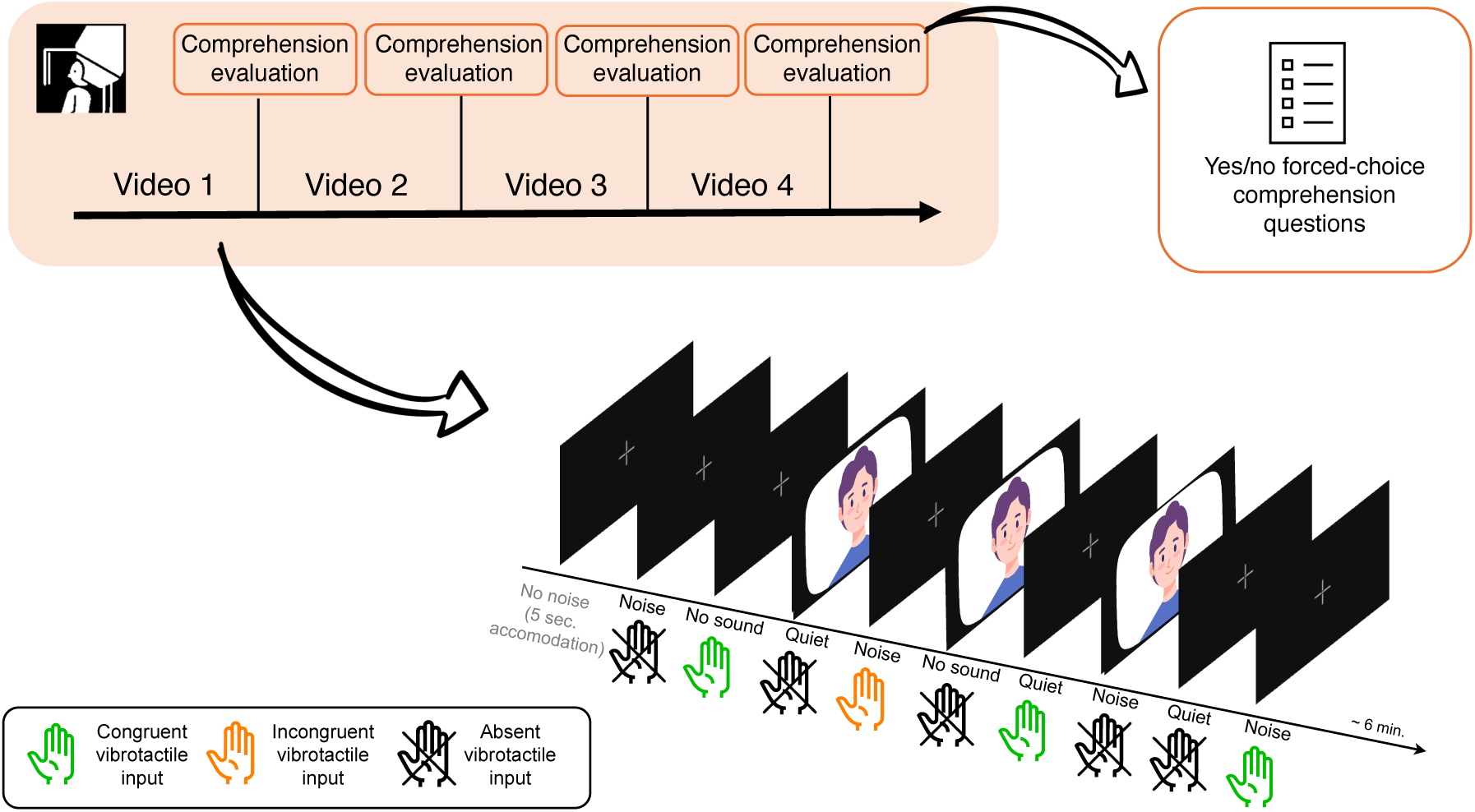
– Illustration of the experiment progression during the MEG acquisition (orange panel) and a time-course of a video stimulus. Participants viewed four videos in total, each segmented into 9 blocks corresponding to the experimental conditions (mean block duration: ∼ 40 s per video). In audio-visual conditions, the visual input was always presented in congruence with the auditory speech signal.

## Results

### Behavioral comprehension evaluation

After watching each of the four videos (Figure 1), participants answered sets of forced-choice yes/no content questions to assess comprehension of the speech material in each experimental condition (i.e., A-only, AV, ATc in quiet; A-only, AV, ATc, ATi in the multi-talker background). Figure 2 displays the distribution of mean comprehension scores, computed as the number of correct answers per condition across all videos. Mean accuracy ranged from 66.7% to 82.5%, suggesting consistent task engagement. Although comprehension in the A-only quiet condition was not at ceiling (79.2 ± 18.4%; mean ± s.d.), performance in all conditions was significantly above the 50% chance level imposed by the yes/no comprehension questions format (all *t*(30) > 3.73, *ps* < .001). The absence of a ceiling effect in quiet likely reflects the complexity of the continuous speech material, which featured information-rich content, and the extensive length of the videos (*Methods: Stimuli*).

**Figure 2.**
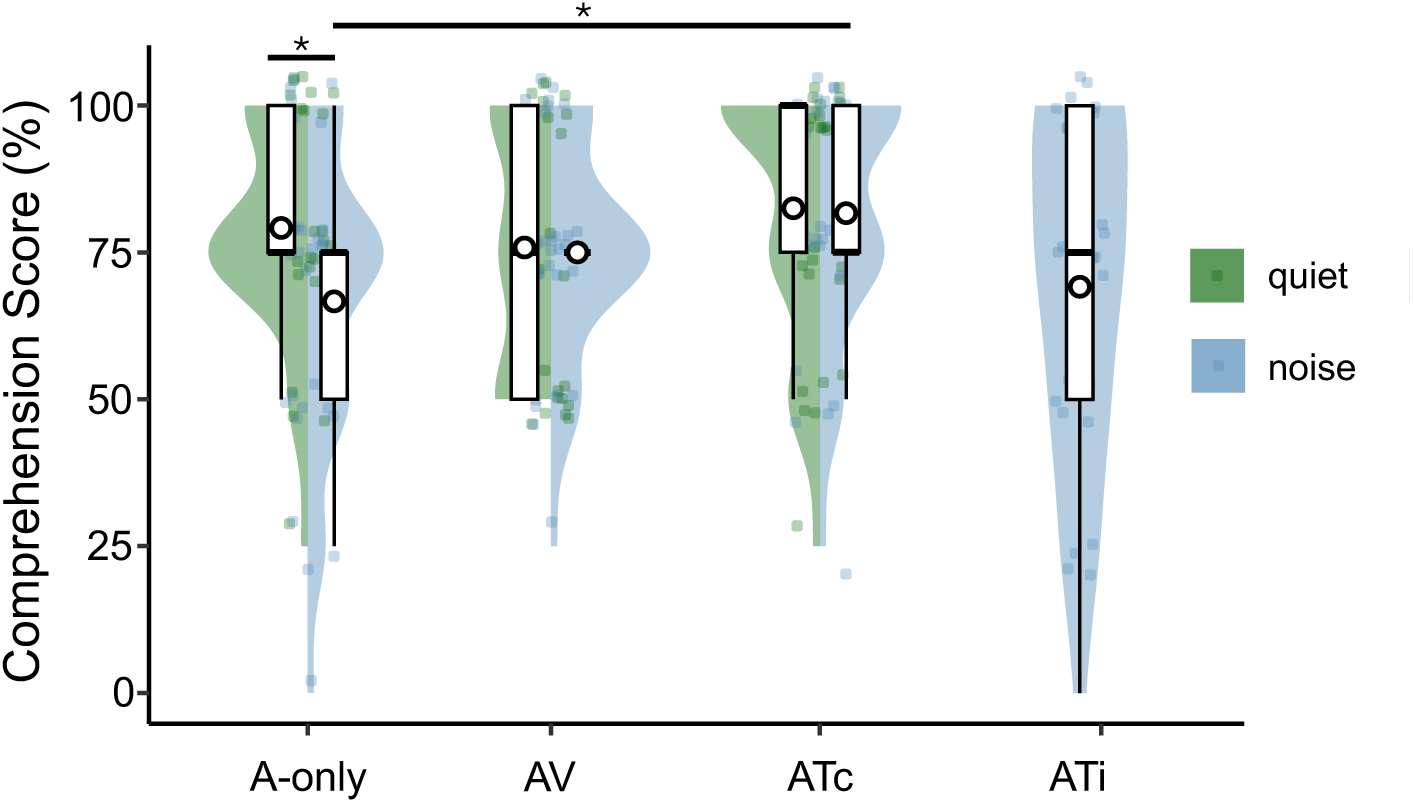
– Comprehension scores (% correct) across experimental conditions. Boxplots represent interquartile range, median (horizontal line) and outliers; white circles indicate group means. A-only: auditory-only, AV: audio-visual, ATc: audio-tactile with congruent vibrations, ATi: audio-tactile with incongruent vibrations. Significance marked for pairwise testing; *p < 0.05.

A logistic mixed-effects regression analysis of the participants’ answers to the content questions revealed that comprehension varied significantly across the experimental conditions (χ²(6) = 13.99, *p* = .029). In quiet, post-hoc pairwise analyses showed that speech comprehension in the A-only condition did not differ from AV (*z* = 0.63, *p* = .98) or ATc (*z* = –0.67, *p* = .52). As expected, the multi-talker background significantly impaired comprehension in the unimodal (i.e., A-only) condition compared to quiet (12.5% decrease; *z* = –2.22, *p* = .039). This deleterious effect of noise was, however, not significant when a supplemental congruent non-auditory sensory input was provided (i.e., quiet *vs.* noise in AV: *z* = –0.15, *p* = .83; quiet *vs*. noise in ATc: *z* = –0.17, *p* = .82). In the multi-talker background, the vibrotactile stimulation significantly improved comprehension compared to the A-only condition when presented congruently (i.e., ATc; 15% improvement, *z* = 2.69, *p* = .016), but not incongruently (i.e., ATi; 3.3% improvement, *z* = 0.57, *p* = .69) with the attended speech. Visual speech, too, provided a modest but not statistically significant comprehension enhancement in noise compared to the A-only condition (8.3% improvement; *z* = 1.45, *p*_unc_ = .073). Notably, the difference between ATc and AV performance was not significant, neither in quiet (6.7% difference; *z* = –1.29, *p* = .98), nor in noise (6.7% difference; *z* = –1.28, *p* = .33). Overall, speech comprehension in quiet was unaffected by the addition of a supplemental, non-auditory congruent sensory signal, whether visual or tactile. Most importantly, while both visual speech and congruent speech-based vibrations mitigated the detrimental effect of the multi-talker background observed in unimodal (A-only) conditions, only the latter led to a significant improvement in comprehension compared to A-only.

### Cortical tracking of speech

Next, we examined the accuracy of the cortical representation of the attended speech envelope (i.e., CTS) across conditions. CTS was quantified using coherence, a well-established measure of CTS^2,31–33^ capturing the phase consistency and amplitude association between two signals on a scale from 0 (no linear association) to 1 (full linear association)^34^. In practice, we computed the coherence between the attended speech temporal envelope and source-localized brain signals, at both syllabic (i.e., 4–8 Hz) and phrasal/sentential (i.e., 0.5 Hz) rates. Given the previously reported hemispheric asymmetries in CTS^2,35^, CTS was computed separately for each hemisphere.

Figure 3 illustrates the cortical distribution of syllabic and phrasal CTS across all experimental conditions. In all cases, CTS sources were present bilaterally in supratemporal auditory cortices (STAC) (see Table 1 for MNI coordinates and statistical significance of the STAC local maxima), confirming the central role of auditory regions in CTS and speech processing. Given our assumption that improvements in speech comprehension in AT conditions stem from CTS refinement in auditory regions, we then analyzed CTS values obtained across conditions at bilateral STAC (Figure 4) using linear mixed-effects modelling, separately for each frequency. All *post-hoc* pairwise statistical comparisons (two-tailed) following these two analyses can be found in Table 2.

**Figure 3.**
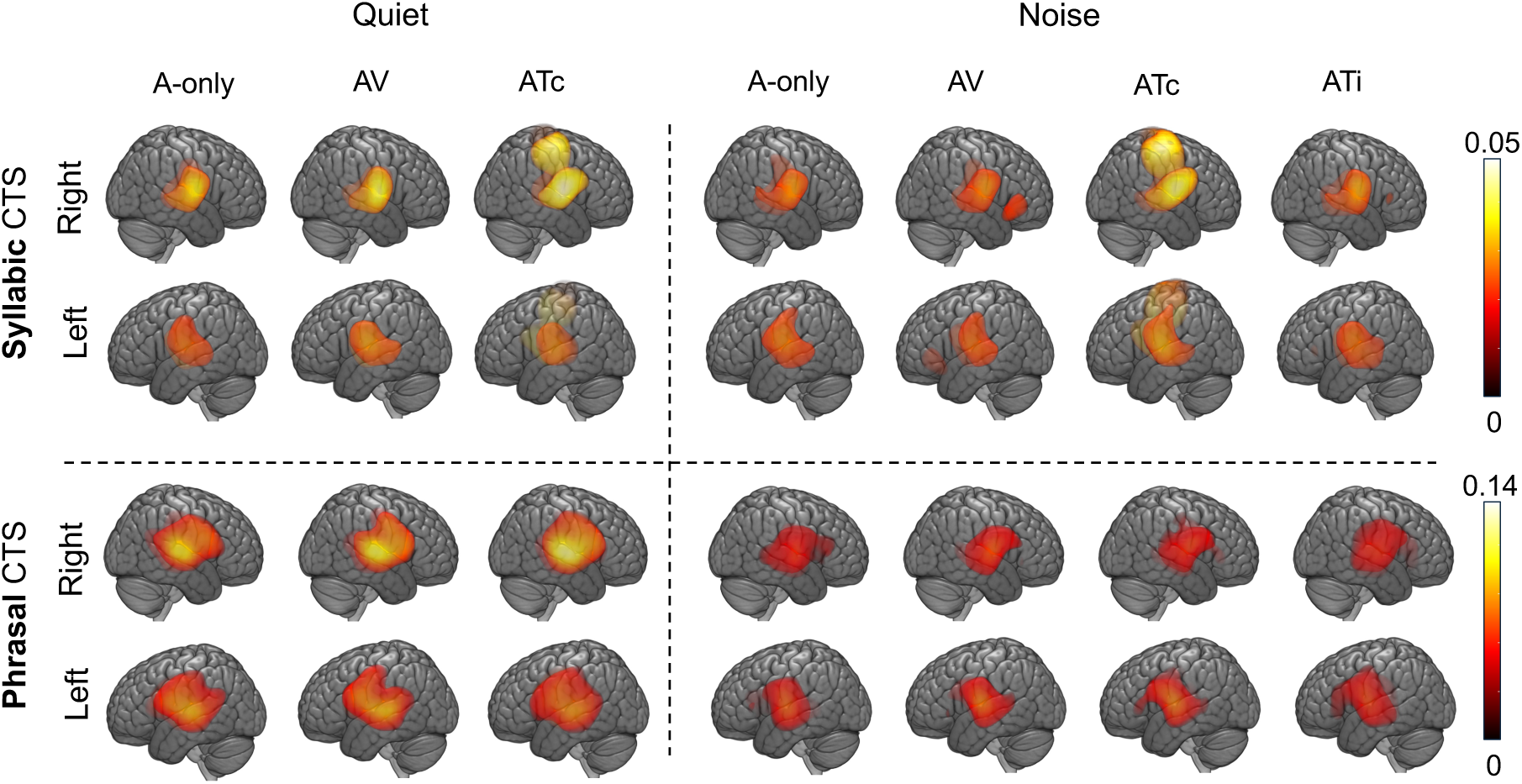
– Brain distribution of significant cortical tracking of speech (CTS) at syllabic (top) and phrasal (bottom) levels, in quiet (left) and with multi-talker background noise (right). Group-level coherence maps (n = 30) obtained by statistically masking above the significance threshold level with the use of nonparametric permutation statistics. A-only: auditory-only, AV: audio-visual, ATc: audio-tactile with congruent vibrations, ATi: audio-tactile with incongruent vibrations.

**Figure 4.**
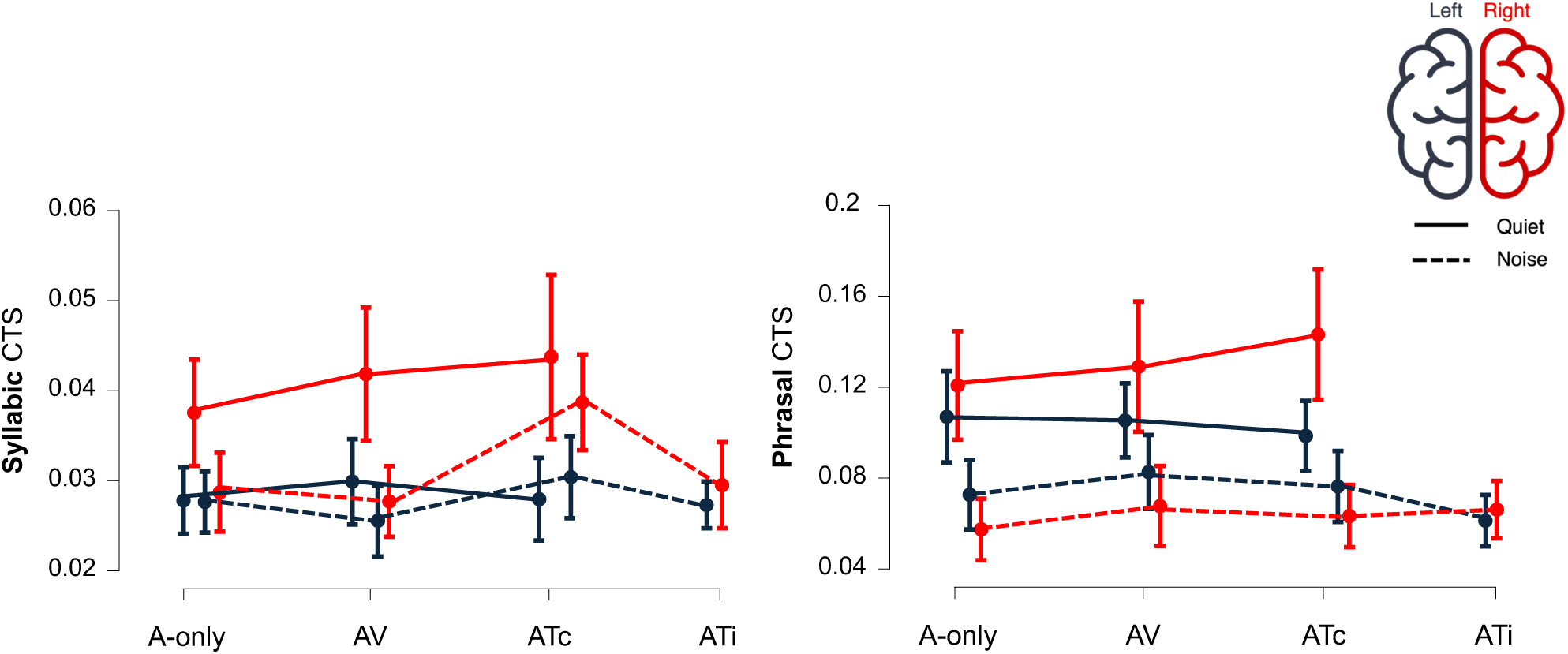
– Impact of experimental condition and hemisphere on the cortical tracking of speech (CTS) at the syllabic (left) and phrasal (right) rates. Means are depicted by the circles and 95% confidence intervals by the vertical lines. Line patterns correspond to the auditory conditions (connected traces – quiet, dashed traces – noise), color indicates the hemisphere in which the CTS was measured (left hemisphere – blue, right hemisphere – red). A-only: auditory-only, AV: audio-visual, ATc: audio-tactile with congruent vibrations, ATi: audio-tactile with incongruent vibrations.

**Table 1.**
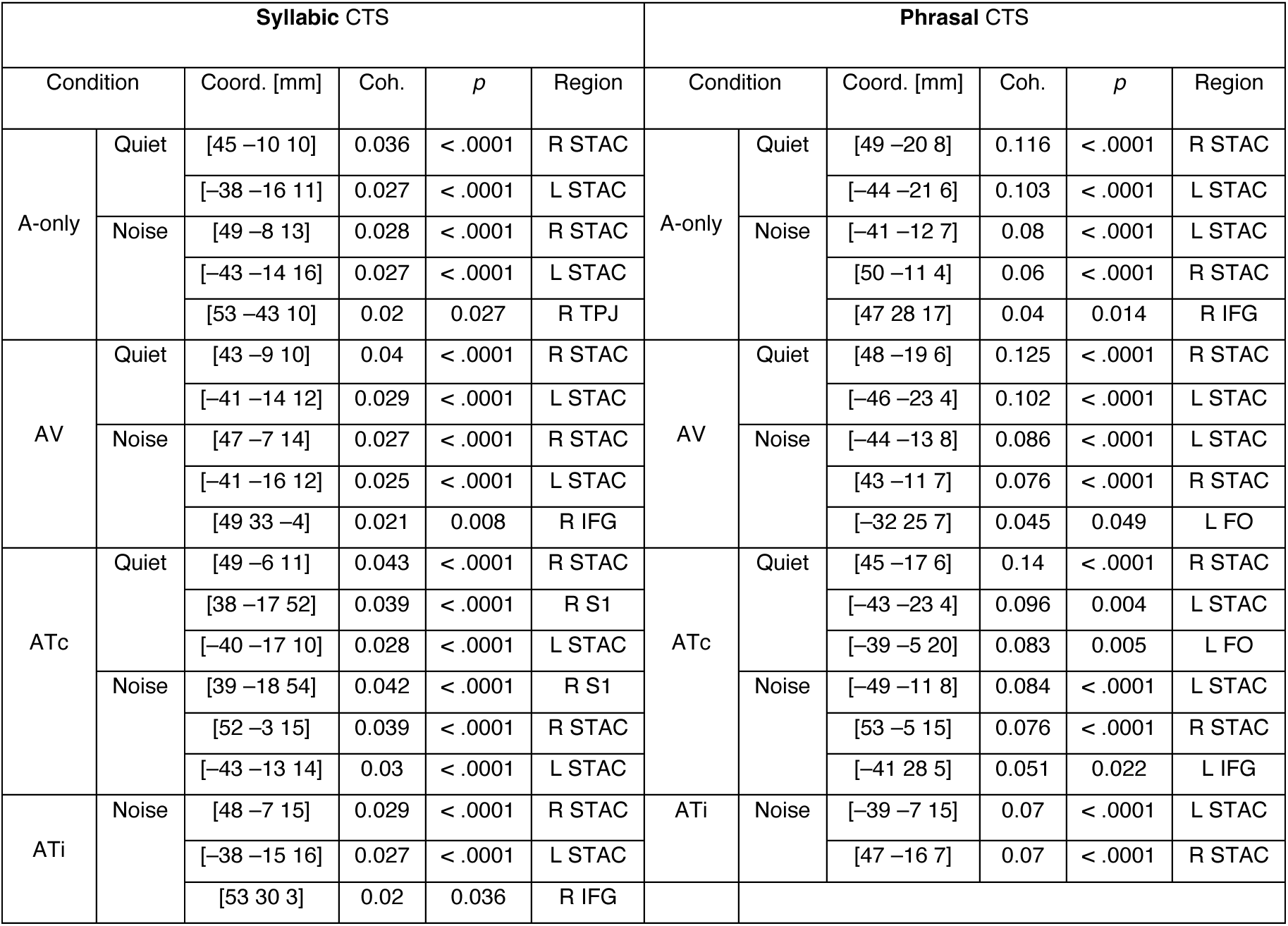
– Significant local maxima of cortical tracking of speech (CTS): peak MNI coordinates [mm], peak coherence values, significance level and anatomical location. A-only: auditory-only, AV: audio-visual, ATc: audio-tactile with congruent vibrations, ATi: audio-tactile with incongruent vibrations, R: right, L: left, STAC: supratemporal auditory cortex, FO: frontal operculum, IFG: inferior frontal gyrus, S1: primary somatosensory cortex, TPJ: temporoparietal junction, Coh.: coherence, Coord.: coordinates.

**Table 2.** – Pairwise comparisons of phrasal and syllabic cortical speech tracking (CTS) values at auditory cortices. Values are reported separately for the left and right hemispheres. P-values (both uncorrected and corrected for multiplicity) reflect pairwise comparisons of estimated marginal means derived from the linear mixed-effects models. A-only: auditory-only, AV: audio-visual, ATc: audio-tactile with congruent vibrations, ATi: audio-tactile with incongruent vibrations, Env.: auditory environment.

| Env. | Hemisphere | Contrast | Syllabic CTS |  |  | Phrasal CTS |  |  |
| --- | --- | --- | --- | --- | --- | --- | --- | --- |
| | | | $t_{403}$ | $p_{uncorrected}$ | $p_{corrected}$ | $t_{403}$ | $p_{uncorrected}$ | $p_{corrected}$ |
| Quiet | Left | A-only vs ATc | -0.210 | 0.834 | 1 | 0.730 | 0.456 | 0.999 |
|  |  | A-only vs AV | -1.161 | 0.246 | 0.987 | -0.146 | 0.884 | 1 |
|  |  | AV vs ATc | 0.951 | 0.342 | 0.998 | 0.876 | 0.381 | 0.999 |
|  | Right | A-only vs ATc | -3.048 | 0.003 | 0.059 | -2.227 | 0.027 | 0.415 |
|  |  | A-only vs AV | -1.518 | 0.129 | 0.902 | -0.543 | 0.587 | 1 |
|  |  | AV vs ATc | -1.530 | 0.127 | 0.897 | -1.684 | 0.093 | 0.817 |
| Noise | Left | A-only vs ATc | -1.041 | 0.298 | 0.995 | -0.386 | 0.699 | 1 |
|  |  | A-only vs ATi | -0.235 | 0.814 | 1 | 0.493 | 0.622 | 1 |
|  |  | A-only vs AV | 0.973 | 0.331 | 0.998 | -0.402 | 0.688 | 1 |
|  |  | ATc vs ATi | 0.806 | 0.421 | 0.999 | 0.879 | 0.379 | 0.999 |
|  |  | AV vs ATc | -2.014 | 0.045 | 0.579 | 0.016 | 0.987 | 1 |
|  | Right | A-only vs ATc | -5.282 | <.0001 | **<.0001** | -0.090 | 0.928 | 1 |
|  |  | A-only vs ATi | -0.642 | 0.522 | 1 | -0.399 | 0.690 | 1 |
|  |  | A-only vs AV | -0.161 | 0.872 | 1 | -0.785 | 0.433 | 0.999 |
|  |  | ATc vs ATi | 4.640 | <.0001 | **0.0001** | -0.308 | 0.758 | 1 |
|  |  | AV vs ATc | -5.120 | <.0001 | **<.0001** | 0.695 | 0.488 | 0.999 |
| Cross-environment | Left | Quiet vs Noise (A-only) | 0.084 | 0.934 | 1 | 2.378 | 0.018 | 0.313 |
|  |  | Quiet vs Noise (ATc) | -0.747 | 0.455 | 0.999 | 1.262 | 0.208 | 0.975 |
|  |  | Quiet vs Noise (AV) | 2.218 | 0.027 | 0.424 | 2.122 | 0.034 | 0.495 |
|  | Right | Quiet vs Noise (A-only) | 3.820 | 0.0002 | **0.004** | 4.806 | <.0001 | **0.0001** |
|  |  | Quiet vs Noise (ATc) | 1.586 | 0.114 | 0.871 | 4.564 | <.0001 | **<.0001** |
|  |  | Quiet vs Noise (AV) | 5.176 | <.0001 | **<.0001** | 6.943 | <.0001 | **0.0002** |

For syllabic CTS, we uncovered a significant main effect of the experimental condition (*F*(6,390) = 11.85, *p* < .0001), of the measured hemisphere (*F*(1,390) = 58.73, *p* < .0001), and a significant interaction between the two (*F*(6,390) = 6.06, *p* < .0001). The main effect of hemisphere indicated that syllabic CTS was overall right-lateralized (left *vs*. right; *t*(403) = –7.54, *p* < .0001), as previously reported ^2^. The hemispheric-specific impact of the multi-talker background revealed a significant CTS decrease in the right hemisphere in both A-only (*p =* .004) and AV (*p* < .0001) conditions. Critically, however, in ATc conditions, syllabic CTS strength remained stable across quiet and noise. In the left hemisphere, multi-talker noise did not impact syllabic CTS in either condition. Additionally, in quiet, no significant syllabic CTS difference between A-only, AV and ATc conditions emerged in either hemisphere. In the multi-talker background, the same pattern emerged in the left hemisphere, but, critically, not in the right, where CTS was enhanced in ATc compared to A-only (*p* < .0001), AV (*p* < .0001) and ATi (*p* <.0001). These results suggest that while a multi-talker background significantly degraded syllabic-level CTS, congruent vibrotactile input (ATc) effectively mitigated this degradation in the right hemisphere.

For phrasal CTS, we observed a significant main effect of the experimental condition (*F*(6,390) = 17.1, *p* < .0001), as well as a significant interaction between condition and hemisphere (*F*(6,390) = 4.25, *p* = .00037), but no significant effect of the hemisphere (*F*(1,390) = 1.9, *p* = .17). In the right hemisphere, the multi-talker noise markedly reduced phrasal CTS compared to quiet in A-only conditions (*p* = .0001). This effect persisted even with visual speech (AV; *p* =.0002) and congruent tactile vibrations (ATc; *p* < .0001). In the left hemisphere, the noise dampening effect did not reach multiple comparison correction significance in any condition. Uncorrected effects were, however, observed in A-only (*p*_unc_ = .018) and AV (*p*_unc_ = .034) conditions, but not in the ATc condition (*p*_unc_ = .21), suggesting that the vibrotactile input may have slightly attenuated the impact of noise at this timescale and hemisphere. Separately for quiet and the multi-talker background conditions, CTS did not differ significantly between A-only, AV, ATc and ATi in either hemisphere. Overall, these results indicate a robust right-hemispheric reduction in phrasal CTS due to the multi-talker background, which was not significantly mitigated by either visual speech or speech-based vibrations.

Cumulatively, these findings confirm the detrimental effect of competing background talkers on both phrasal and syllabic CTS of the attended speaker at the STAC. Chiefly, they underscore the key enhancement of syllabic CTS at the right STAC induced by congruent speech-derived vibrotactile stimulation in the multi-talker background.

### Linking audio-tactile CTS enhancement to speech comprehension

After uncovering significant audio-tactile enhancing effects in both behavioral and CTS measures, we aimed to investigate whether these two effects were linked. In practice, we tested whether the right STAC syllabic CTS increase in the ATc multi-talker background (i.e., the difference between A-only and ATc, AT–A-only) correlated positively with speech comprehension performance (percentage of correct responses to comprehension questions in the ATc multi-talker condition). This analysis revealed a significant positive correlation between the two (Figure 5, τ(28) = 0.27, *p* = .03). Thus, the syllabic CTS enhancement provided by congruent speech vibrations in the right auditory cortex was linked to increased speech understanding, suggesting a brain-behavior association.

**Figure 5.**
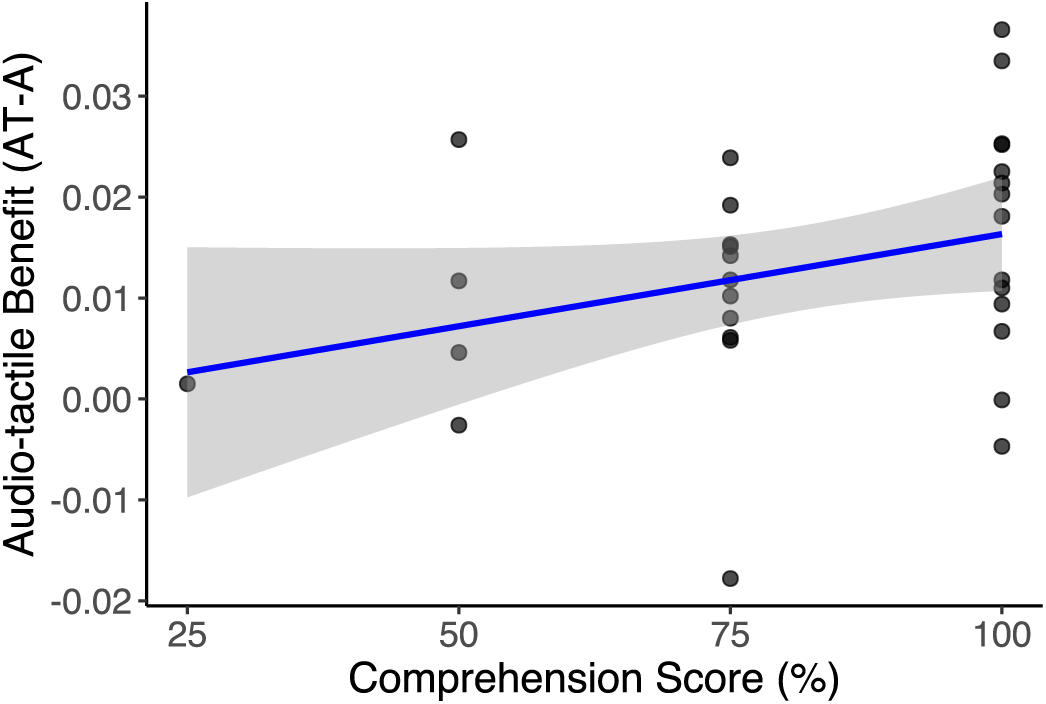
– Behavioral relevance of audio-tactile (AT) cortical tracking of speech (CTS) enhancement. Correlation between the syllabic CTS increase in right-hemispheric supratemporal auditory cortex in the ATc condition (relative to the auditory-only condition) and comprehension score in the same condition.

### Extra-auditory CTS sources

Several loci exhibiting significant CTS outside STAC, previously implicated in CTS^2,17,32^, were observed, particularly in noise conditions (Table 1). Their emergence is most likely due to a noise-dependent reduction of syllabic and phrasal CTS at STAC, enabling weaker CTS sources to attain significance during permutation testing (*Methods: Statistical Analyses)*. More importantly, however, in ATc conditions (both in quiet and multi-talker noise), we found a significant syllabic CTS source in the right primary somatosensory (S1) cortex (Figure 3), which likely reflected the direct somatosensory processing of the left-hand vibrotactile stimulation. Given this, we next examined if individual differences in this cortical-level somatosensory processing were related to the observed AT CTS benefit using correlational analyses. More specifically, we sought to uncover whether the latter was driven by the processing of the vibrotactile input in right S1 cortex— potentially reflecting better intrinsic sensory encoding—rather than being STAC-specific. Results indicated no association between syllabic CTS at S1 cortex and either the ATc syllabic CTS benefit (*r*(28) = 0.09, *p* = .31) or the ATc syllabic CTS values (*r*(28) = 0.17, *p* = .19). As such, these findings suggest that the observed CTS benefit was not driven by a somatosensory encoding-related mechanism but rather resulted from an auditory-specific CTS enhancement.

### Functional connectivity changes linked to audio-tactile CTS enhancement

Next, we examined if the significant syllabic CTS enhancement in the multi-talker ATc condition (vs. A-only) was accompanied by broader functional connectivity changes of the right STAC. We hypothesized that, in the ATc condition, this area synchronized more strongly with associative cortical areas involved in AT multisensory integration^36^, or with somatosensory cortices (as formerly shown with visual cortices in AV speech^37^). To address this, we conducted a seed-based functional connectivity analysis at the right STAC local maxima, comparing multi-talker ATc and A-only conditions. Functional connectivity was quantified using the phase locking value (PLV), a phase alignment measure ranging from 0 (no synchronization) to 1 (full synchronization)^38^. This analysis covered both syllabic and phrasal frequency bands, as well as three additional frequency bands (i.e., alpha: 8–12 Hz, low-beta: 12–21 Hz and high-beta: 21–30 Hz) known to be involved in speech processing^39,40^. Although AT CTS effects were theta-band restricted, changes in STAC’s synchronization with other brain areas at non-theta frequencies could facilitate the temporal coordination of neural activity within STAC, optimizing CTS^41^.

Results (Figure 6) revealed that the addition of the congruent speech-based vibrotactile stimulation did not influence right STAC functional connectivity with somatosensory-specific regions in any frequency band. It was, however, associated with an increased low beta-band functional connectivity with the right angular gyrus (AG: [51, –64, 32] mm; *p* = .049, permutation test, two-tailed, *n*_perm_ = 2000), as well as with the posterior part of the right inferior temporal gyrus (ITG: [46, –51, –14] mm; *p* = .0495, permutation test, two-tailed, *n*_perm_ = 2000). Additionally, we observed a significant decrease in alpha-band connectivity with the precuneus ([11, –46, 59] mm; *p* = .0455, permutation test, two-tailed, *n*_perm_ = 2000). A secondary analysis seeded from the left STAC did not uncover any significant functional connectivity differences between conditions.

**Figure 6.**
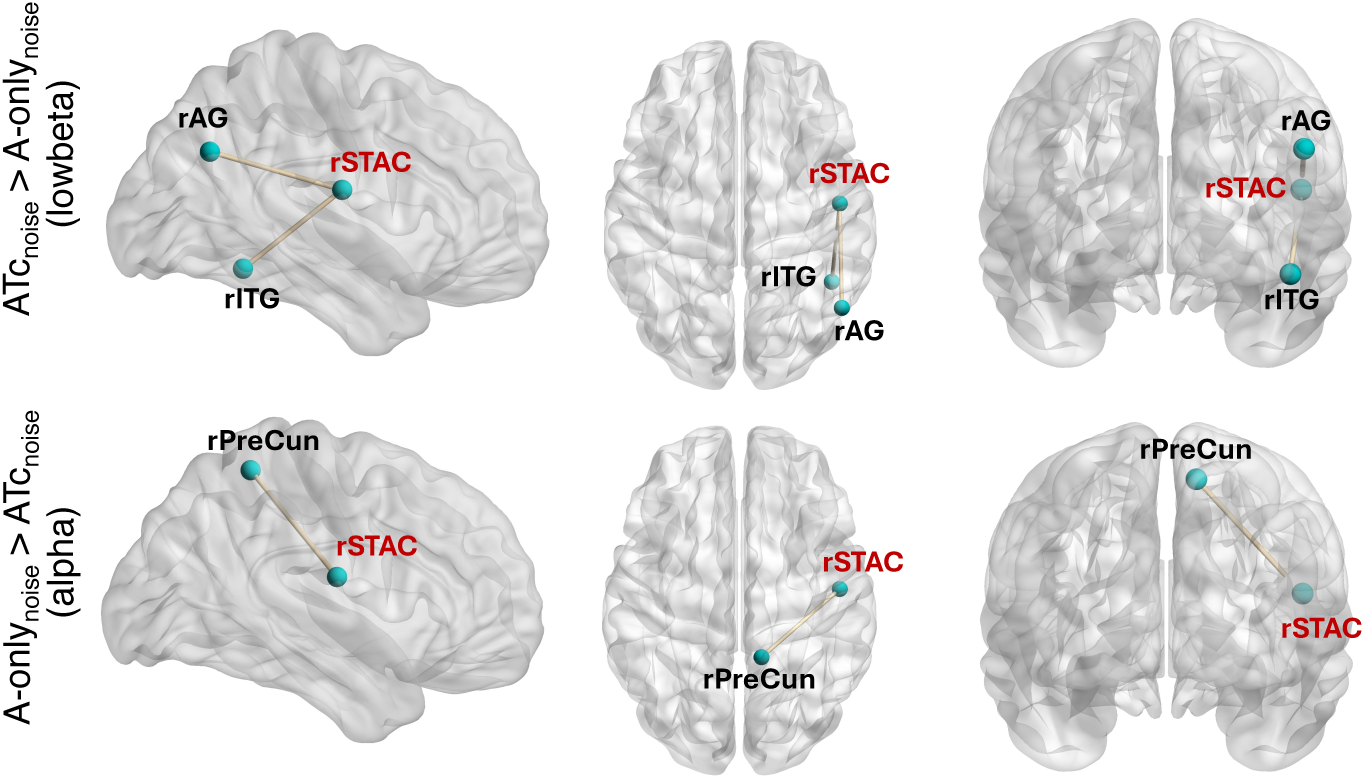
– Significant functional connectivity changes in the audio-tactile multi-talker condition compared to the unimodal, auditory-only condition. Functional connectivity changes seeded from CTS local maximum in right supratemporal auditory cortex (STAC, in red). r = right hemisphere, AG = angular gyrus, ITG = inferior temporal gyrus, PreCun = precuneus, A-only = auditory-only, ATc = audio-tactile with congruent vibrations. Figures created using BrainNet viewer^97^.

Overall, these findings indicate that in the ATc compared with the A-only condition, right STAC functional connectivity is increased specifically with higher-order, ipsilateral, associative parietal and temporal neocortical regions, rather than with somatosensory-specific areas, potentially reflecting enhanced supramodal integration mechanisms.

## Discussion

This study demonstrates that, in a complex multi-talker auditory scene, the concurrent presentation of congruent speech-derived vibrotactile stimulation is associated with increased CTS of attended speech syllabic rhythms in the right STAC. Behaviorally, comprehension in this audio-tactile condition also improved relative to A-only, and, importantly, positively correlated with the extent of the AT CTS increase. Of note, this effect was accompanied by frequency-specific changes in functional connectivity between the right STAC and ipsilateral, higher-order associative parieto-temporal neocortical areas, suggesting the complementary emergence of a broader, long-range network-level modulation during audio-tactile speech processing.

### Vibrotactile enhancement of speech comprehension

Behaviorally, comprehension of the attended speaker in the multi-talker background improved significantly (15%) when listeners received speech-derived vibrotactile stimulation congruent with the attended speech stream. These results align with previous reports of behavioral, sentence-level vibrotactile comprehension enhancement^6,28^, further highlighting that this multisensory effect can emerge even during naturalistic, connected speech listening. In quiet, this enhancement was markedly reduced (3.3%) and non-significant, indicating its contingency on the presence of background noise. This noise-specific effect conforms to the inverse effectiveness principle of multisensory integration, which states that as response accuracy to unimodal stimuli decreases, the strength of multisensory integration increases^42^. Incongruent vibrations also did not yield a significant benefit compared to A-only, underscoring the added essential role of synchronicity between sensory modalities (i.e., temporal rule of multisensory integration)^43^. This corroborates previous observations showing significant tactile speech enhancement with congruent, but not incongruent, vibrotactile speech input^19^. Importantly, the emergence of this benefit in untrained listeners with no prior exposure to the vibrotactile stimulation reinforces theories that conceptualize speech perception as a multimodal sensory phenomenon, with modality-invariant speech cues aiding comprehension regardless of the transmission channel^44–46^.

### Vibrotactile enhancement of cortical tracking of speech and impact on behavior

In A-only, we observed a reduction in both syllabic and phrasal CTS in the presence of the multi-talker background, with a stronger dampening in the right compared to the left hemisphere, consistent with previous studies^2,35^. Under these adverse auditory conditions, congruent speech-derived vibrotactile stimulation significantly restored syllabic CTS at the right STAC, but not phrasal CTS. The syllabic-level specificity of this effect may reflect intrinsic processing constraints of the somatosensory system, as somatosensory (S1 and S2) cortices operate within short temporal integration windows (i.e., <200ms)^47^. Phrasal-level information would require a longer temporal window for effective integration (up to 1–2 seconds), likely exceeding the somatosensory cortex’s typical range. This was confirmed here by the consistent presence of an S1 cortex source in ATc conditions for syllabic, but not phrasal, CTS (Figure 3; Table 1). The right-hemispheric lateralization of this effect may further stem from (i) the established role of right auditory areas in syllabic-level processing and CTS^48,49^, (ii) robust audio-tactile integrative effects observed in the hemisphere contralateral to the side stimulated tactilely (left-hand stimulation in our study)^50^, and (iii) the right hemisphere’s stronger engagement in audio-tactile integration and multisensory processing^51–53^. Nonetheless, whether this right-hemispheric CTS enhancement would be maintained even when providing vibrotactile stimulation to the right palm remains an open question.

Another key finding of the current study was that, in multi-talker conditions, the greater the syllabic CTS increase at right STAC in audio-tactile compared to the unimodal condition, the better the participants’ performance on questions related to the speech content. This brain-behavior association substantiates previous findings of a positive link between audio-tactile CTS and speech intelligibility^27^. In addition, it reinforces the central role of syllabic CTS in supporting speech understanding in noisy environments, as demonstrated using theta-band vibrotactile input^28^, transcranial current stimulation^54^, and in clinical or developmental populations with reduced speech-in-noise ability^17,32,55^. Taken together, these results suggest that syllabic CTS may serve as the neural scaffold for audio-tactile comprehension enhancement in multi-talker conditions.

### Functional connectivity changes underlying audio-tactile CTS enhancement

In the ATc multi-talker condition, we evidenced increased ipsilateral low-beta functional connectivity between the right STAC and the right AG and posterior ITG. The AG is a well-known convergence zone of multisensory input^56^, critical in binding cross-modal information for conceptual representation. Similarly, the right ITG is a high-level multimodal area involved in cross-modal expectation violation^57^. Given the added role of beta-band oscillations in top-down predictive coding during speech processing^58^, this increased synchronization may suggest a top-down mechanism of contextual filtering through which STAC auditory representations that mismatch the concurrent tactile input are suppressed selectively, enhancing the accuracy of syllabic CTS. Furthermore, while lexico-semantic processing is typically left-lateralized, right-hemispheric AG and ITG have both been linked to such processes^56,59,60^. Thus, these regions may further refine the temporal resolution of speech envelope encoding at STAC through lexico-semantic priors, using the perceived audio-tactile speech cues to constrain linguistic predictions, facilitating attended speech stream segmentation. Given the absence of left-hemispheric involvement, however, this interpretation warrants further investigation, for example through manipulations of tactile laterality and speech input lexico-semantic predictability. Notably, these mechanisms may work in parallel, jointly enhancing syllabic CTS.

Decreased alpha-band functional connectivity between the right STAC and the right precuneus was also observed in the multi-talker ATc condition (*vs*. A-only). The precuneus, a key node of the default-mode network (DMN), is involved in internally-oriented processing^61^, while alpha oscillations play a key role in mediating DMN functional connectivity^62^ and disengagement from the sensory environment^63^. This decrease might thus reflect a shift toward externally driven processing in the ATc condition, contributing to better CTS by freeing up neural resources for speech-related (multi-)sensory processing.

These findings highlight a broader, long-range network-level mechanism during ATc, where higher-order, multisensory areas interact with STAC cortical dynamics, potentially refining speech envelope tracking through beta-mediated predictive feedback and alpha-mediated attentional shifts. This underscores the engagement of associative cortical regions—rather than direct somatosensory-auditory connections—under adverse auditory conditions with audio-tactile multisensory input.

### Comparison with visual enhancement

We found no significant behavioral difference between the benefit provided by the congruent vibrotactile stimulation and the 8.3% benefit provided by visual speech (i.e., seeing the speaker’s articulatory gestures). This AV enhancement, however, did not translate to enhanced CTS, diverging from prior studies showing increased speech understanding and CTS in multi-talker conditions with visual speech support^4,16,17^. This might be due to the employed SNR (i.e., 0 dB), as SNRs that maximize AV perceptual benefit have been found to be much lower (i.e. –9 dB)^18^. In studies which uncovered AV effects at higher SNRs^17^, the used visual stimulus material could underlie the discrepancy in the uncovered effects, given that the narrators in the present study exhibited minimal facial movement and expression. Moreover, participants were not explicitly instructed to focus on the narrators’ lips, which are essential for encoding envelope information^64^. AV and AT CTS enhancement may rely on similar mechanisms only if both provide the same supramodal category of speech input (here, amplitude speech modulations). Insufficient visual attendance to the speaker’s lips, further disrupted by the speakers’ eye movements due to teleprompter reading, may have limited the extraction of articulatory cues, thereby preventing speech envelope encoding in auditory cortices^64,65^. In contrast, tactile input in AT conditions is often processed in a bottom-up manner, exerting modulatory effects on neural activity even in the absence of measurable behavioral effects^27,66^. As such, tactile speech cues may have had a more pronounced effect on CTS responses. Alternatively, given the weak direct correlation between lip movements and speech envelope^31^, the tactile vibrations—directly derived from amplitude modulations—were a more robust transmission channel for envelope information. We underscore the importance of more careful gaze behavior control in future studies contrasting AV and AT speech enhancement.

### Limitations

The uncovered vibrotactile syllabic CTS enhancement is suggestive of an integrative mechanism, whereby neural populations in the STAC exhibit a stronger response in ATc relative to the response to unisensory stimuli (auditory speech and speech-derived vibrations) alone. A limitation of the current study is that we cannot resolve whether this effect’s underlying computation is additive (i.e., a linear summation of unimodal responses; AT = A-only + T-only) or superadditive (i.e., the summation exceeding the linear sum of unimodal responses; AT > A-only + T-only)^42^. Although the experimental design (*Methods: Stimuli*) included unimodal, T-only and V-only conditions (not described here), the computation of CTS using coherence analyses prevents a straightforward computation of this aspect. As coherence is a bounded and positively biased measure, T-only CTS values at STAC, while positive, may be non-significant and not reflect genuine speech-brain coupling. Using them for additive predictions (A-only + T-only) could artificially inflate the obtained sum, concealing any potential superadditive effects. Alternative CTS metrics, such as cross-validated encoding/decoding models, could potentially better uncover the precise nature of these audio-tactile multisensory effects, already reported to exhibit superadditive properties^27^ (but see^26^ for diverging results).

An added limitation of this work is the use of yes/no content questions for the comprehension evaluation. While suitable for the employed material (i.e., connected speech), these lack granularity and are more sensitive to random guessing than more in-depth comprehension measures. This could have contributed to the absence of behavioral AV effects, widely documented previously. In keeping with the ecological validity of the present design, future studies could complement this evaluation with more complex comprehension tasks, such as free recall or paraphrasing. Alternatively, measures such as sentence recognition thresholds could provide greater sensitivity in evaluating comprehension, but this would diminish the ecological validity of the experimental design.

### Conclusion

This study demonstrates that, in a multi-talker scenario, supplemental, congruent speech-derived vibrotactile stimulation enhances comprehension of an attended speaker and gives rise to two distinct neural manifestations: improved temporal envelope encoding at the right auditory cortex (contralateral to the stimulated side), and changes in functional connectivity between this auditory area and ipsilateral, higher-order multisensory regions. These findings shed light on the neural pathways involved in audio-tactile speech enhancement, highlighting the potential of speech-derived vibrotactile stimulation to modulate speech-related neural activity towards improved speech comprehension in adverse auditory conditions.

## Materials and methods

### 1. Participants

Thirty native French-speaking adults (15 female) aged 25.5 ± 5.0 years (mean ± s.d.) participated in this study. All participants were right-handed based on the Edinburgh Handedness Inventory^67^. All had normal hearing as indicated by hearing thresholds (air-conducted) of ≤25 dB HL (hearing level) obtained during pure-tone audiometric testing at all standard frequencies between 125 Hz and 8000 Hz. Participants’ central auditory perception was also evaluated using a standardized French language battery^68^ and a speech-in-noise recognition threshold measurement using the French Sentence Test for Speech Intelligibility in Noise (FIST)^69^ (procedure described in Răutu et al.^6^) with no participants demonstrating atypical auditory abilities. They all had normal or corrected-to-normal vision and normal somatosensory perception, with the latter being confirmed through a vibrotactile threshold assessment ^6^. Participants completed a brief questionnaire to report their current medication or history of psychiatric, neurological, auditory or somatosensory disorders. No relevant pathology was indicated. The study was approved by the ULB—Hôpital Erasme Ethical Committee (P2012/049). Written informed consent was obtained from all participants prior to testing and they received monetary compensation for their participation.

### 2. Stimuli

Stimuli were derived from eight audio-visual recordings of trained, native French-speaking narrators (one male and one female, four recordings per narrator) speaking for ∼6 minutes (6.0 ± 0.3 min). During video recording, speakers were instructed to speak with little to no facial expressions and head or body movements. The linguistic content of the recordings included the characteristics and habitat of eight uncommon animals: the axolotl, tardigrade, dugong, kiwi, wombat, Dumbo octopus, Trionyx and aye-aye. This specific topic was chosen for its emotional neutrality^70^, richness in informational content, and, given the animals’ rarity, the difficulty to intuit information during the comprehension evaluation. This was confirmed *a posteriori* following the MEG data recording, as no participant indicated in-depth familiarity with the heard information. Speech content comprehension was assessed using yes/no forced-choice questions, targeting condition-specific informational content present only in the attended speech stream. This approach was chosen because of the continuous and naturalistic nature of the experiment, ensuring that participants engaged with the narrative without disruptive task switching (e.g., target-word detection or button press responses). Videos were first edited using Final Cut Pro (v10.6.5, Apple Inc.) to remove prolonged pauses and errors. Recordings were slowed down as needed to ensure clear comprehension and consistent syllabic rhythm across recordings (final mean syllabic rate: 5.61 ± 0.28 Hz). Based on the recordings, experimental stimuli were prepared as described below.

#### Auditory stimuli

Video soundtracks were extracted, denoised and intensity-normalized across videos (root mean square procedure) using the Audacity audio editing software (v3.0.2; https://www.audacityteam.org). For the multi-talker conditions, an auditory background composed of 6 native French speakers (3 females) talking simultaneously^71^ was normalized and mixed with the soundtracks at 0 dB SNR using custom Matlab (R2021a, TheMathWorks) scripts.

#### Tactile stimuli

Tactile stimuli consisted in vibrations derived from the speech temporal envelope of the unaltered (i.e., unmixed with the multi-talker background) speech signal of each video. The generation of the vibrations was done using custom Matlab (R2021a, TheMathWorks) scripts^6^. First, for each normalized clean audio track from the original 8 videos, the corresponding speech envelopes were extracted using an established approach^17^ which entailed passing each audio track through a 31-channel gammatone filter bank ranging from 150 to 7000 Hz (Mel scale), followed by computing the temporal envelope of each channel using the Hilbert transform. Then, all channel-specific envelopes were averaged to obtain a composite speech temporal envelope. The gammatone filter frequency interval was chosen to overlap with both the average speech spectrum and the frequency range of human auditory perception^72^. Figure 7A illustrates the envelope obtained from an excerpt of one audio segment. Finally, the computed temporal envelope was used to amplitude modulate a 150 Hz sinusoidal carrier, the prime sensitivity range of the principal vibratory mechanoreceptors, Pacinian corpuscles^73^.

**Figure 7.**
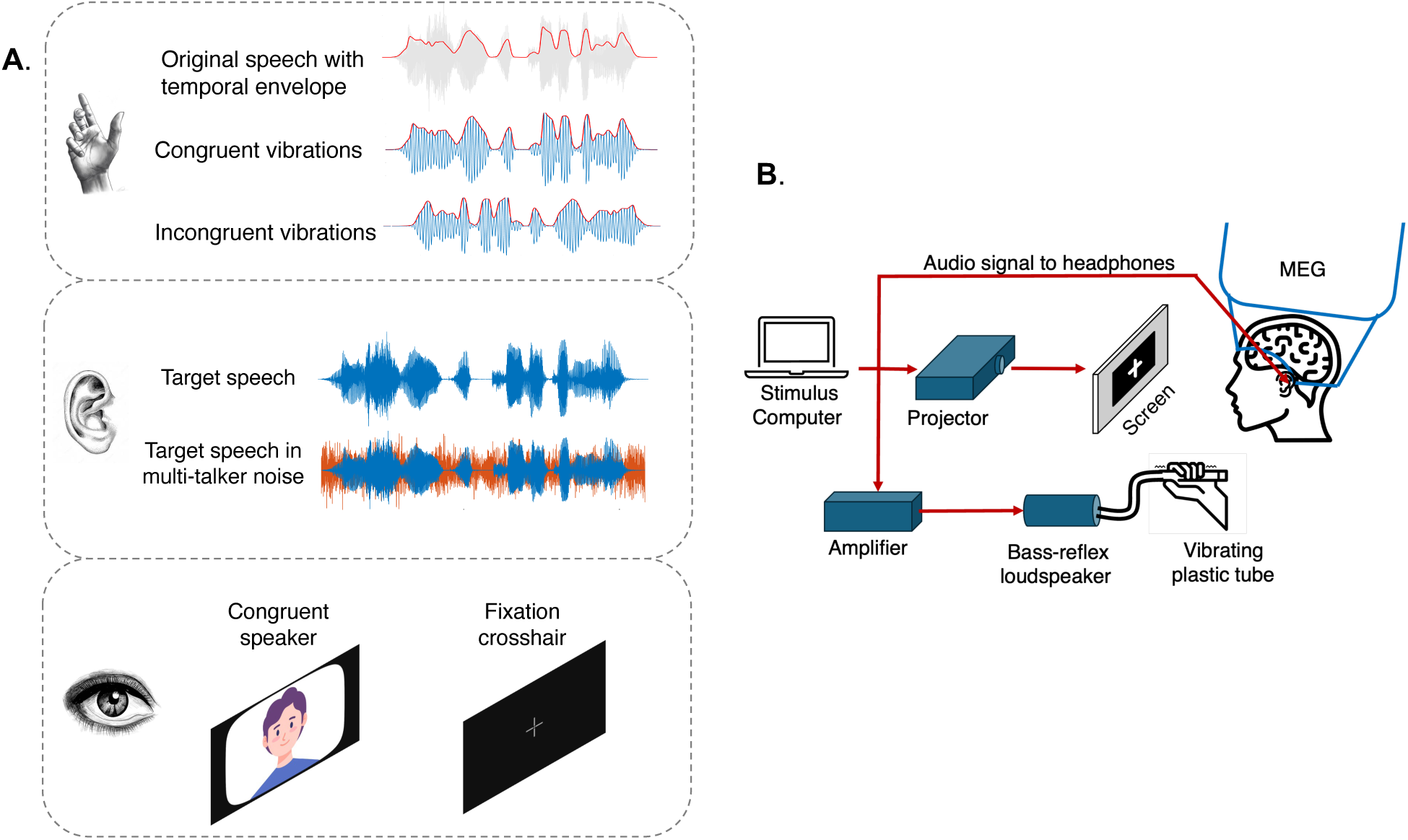
– Experimental material and setup. (A) Vibrotactile inputs (palm symbol) were derived from the speech envelope of the heard speech signal (first row, in red); congruent vibrations (middle row) had amplitude variations (in red) as per the speech temporal envelope, while incongruent vibrations (bottom row) also had envelope-based amplitude variations (in red) but not aligned with the heard speech. Auditory (ear symbol) inputs consisted in clear speech or speech mixed with multi-talker noise (in orange). Visual (eye symbol) information was either the speaker talking or a fixation crosshair. (B) A main stimulus computer was used to simultaneously deliver the vibrotactile, video and auditory stimulation. Auditory signals were transmitted through MEG-compatible earphones. Visual stimuli were projected on a MEG-compatible screen through a dedicated projector. Vibrotactile signals were transmitted through an amplifier and a bass-reflex speaker, reached participants’ palms through a plastic tube with a vibrating silicone ending.

#### Visual stimuli

Depending on the experimental condition, the visual stimuli consisted of either the original video of the speaker or a fixation cross. Minimal editing was applied to the visual component of the speaker recordings, comprising only color and luminance equilibration across videos, alignment of the faces of the speakers and centering them in the frame and the addition of a black border (all editing done in Final Cut Pro, Apple Inc.). For non-visual conditions, a central grey crosshair was presented against a black background.

#### Final video generation

Final video editing and condition assignment was done using custom Python scripts (v3.9.5, Python Software Foundation, https://www.python.org/). Each of the original recordings was divided into nine equal blocks, which were assigned to nine experimental conditions. Of these, seven blocks included sound and two were soundless (i.e., V-only and T-only, not analyzed in the current paper). To evaluate the role of the sensory modality on speech comprehension and CTS, the blocks with auditory content were either unimodal (A-only) or bimodal; for the latter, the auditory input was paired with either visual (i.e., AV) or tactile (i.e., AT) input. For each sensory input type (A, AV, or AT), two blocks were presented: one in which the attended speech was presented in quiet, and the other in which it was embedded in multi-talker background, totaling 6 conditions. For the 4 multisensory conditions included in these 6, the visual/tactile signal was always synchronous with the attended speech stream. To assess the effect of vibration congruency on the presumed effect of vibrotactile stimulation on CTS enhancement in multi-talker noise, a 7^th^ condition presented vibrations derived from the speech envelope of a distinct audio segment from the same recording (i.e, ATi; Figure 7A).

Figure 1 illustrates an example of a video used during the experiment. The first 5 s of each video featured the original audio of the attended speaker, enabling participants to correctly identify the narrator’s voice. The assignment of conditions to the 9 blocks was pseudo-randomized, with the first condition always containing sound (for continuity with the initial 5 s of the narrator voice), noise conditions always alternating with either quiet or soundless conditions, and successive conditions always presenting differing sensory input types (for example, two AT conditions never followed one another). Based on these constraints, 4 condition orders were established and for each of the original 8 videos, four versions were created corresponding to each of these orders. Video-order assignment was counterbalanced across participants, and we monitored the frequency of the presentation order of the video (e.g., how often a specific video was shown first) and of the condition order (e.g., how frequently the 1^st^ video presented the conditions in the 1^st^ order). 4 videos were presented to each participant, resulting in a data length of ∼2.5 min per condition. Each of the 4 presented videos had a distinct condition order, and the male and female narrators were alternated (2 for each).

### 3. Procedure

The experimental stimuli presentation and neural data recording took place in a quiet, magnetically shielded room^74^. Participants were seated comfortably in the MEG chair with their head inside the MEG helmet. Their arms were placed in front of them, and they were asked to hold with their left hand a vibrotactile stimulation tube, without exerting force. The left hand was preferred over the right as stronger cortical facilitation effects in primary and secondary contralateral somatosensory cortex have been linked to left-handed vibrotactile stimulation in right-handers^75^. Moreover, no significant left-right hand difference has been found in the effects of vibrotactile stimulation on speech-in-noise understanding^6^. Prior to recording, participants were instructed to listen to the narrator’s voice as attentively as possible even in the presence of any background noise, to watch the videos when presented (otherwise to focus on the fixation cross), to maintain contact with the vibrating tube continuously throughout the video, and to move as little as possible. After each MEG recording (one per watched video), participants were asked 9 yes/no comprehension questions corresponding to the 9 experimental conditions from the video. Following the videos, a 5-min MEG recording was conducted with participants at rest (eyes open, no audio, crosshair displayed).

### 4. Hardware

The experimental setup is illustrated in Figure 7B. Recordings were played from the stimuli presentation computer using VLC media player (v3.0.12, VideoLAN Project, GNU General Public License). Videos were displayed on a back-projection screen placed ∼1 m in front of the participants. Auditory stimuli were transmitted binaurally via MEG-compatible earphones (TIP 300, Natus Inc., Wisconsin, USA) with noise-reducing inserts (ER3-14A, Etymotic Inc., IL, USA), with an average sound intensity of ∼65 dB SPL (sound pressure level) as assessed at the earphone level with a sound level meter (Sphynx Audio System). The vibrations were transmitted using a MEG-compatible vibrotactile stimulator similar to the one used in Caetano and Jousmäki^23^ and other studies^6^. Vibrotactile input was sent from the experimental computer to an amplifier connected to a bass-reflex loudspeaker, which then delivered the vibrations through a blind-ended rigid plastic tube (⌀38 mm, 3-m long). The ending of the plastic tube was made from a flexible, vibratory-purposed silicone that had sound-attenuating material (cotton wool) at the closed end. This limited the vibrating tube area to a palm-sized region, whose limits were visually marked. Participants were instructed to respect the limits of these markings while in contact with the tube. The tube was partially fixed laterally to the participants such that it could be held passively. Amplifier settings were kept constant throughout the experiment, ensuring a fixed vibration intensity across participants. The vibration magnitude was chosen to elicit a clear sensation of vibration during stimulation (as evaluated during behavioral piloting, *n* = 5), without the concomitant emergence of any auditory percept. This intensity was reduced by approximately 10 dB during vibrotactile threshold measurements.

### 5. MEG data acquisition

MEG signals were recorded (sampling rate, 1 kHz; band-pass filter, 0.1-330 Hz) at the HUB—Hôpital Erasme using a whole-head MEG system (Triux, MEGIN Oy, Espoo, Finland). The sensor array of the MEG system comprised 306 sensors grouped in 102 triplets of one magnetometer and two orthogonal planar gradiometers. The MEG system was placed in a lightweight magnetically shielded room^74^. The audio presented to the participants was simultaneously recorded (sampling rate, 1 KHz; low-pass, 330 Hz) with the MEG signals to synchronize MEG and audio signals during the data analysis. To continuously monitor the participants’ head position within the MEG helmet, four head position indicator (HPI) coils were attached to their scalp prior to the start of the MEG session. The position of the coils with respect to three anatomical fiducials (nasion and the left/right preauricular points), as well as to at least 300 points on the surface of the head were digitized with an electromagnetic tracker (FASTRAK, Polhemus, Colchester, USA). Following MEG, participants underwent a high-resolution 3D cerebral magnetic resonance imaging (MRI) T1-weighted scan in a hybrid 3T PET-MR scanner (SIGNA, GE Healthcare, Wisconsin, USA).

### 6. MEG data analysis

#### MEG Preprocessing

First, to suppress external sources of magnetic interference and correct for any head movement, MEG data was preprocessed off-line using the temporal signal space separation (tSSS) algorithm implemented in MaxFilter (Neuromag, Elekta; correlation limit 0.9, segment length 10 s)^76,77^. Then, to eliminate physiological artifacts, independent component analysis^78^ using the FastICA algorithm was ran and 30 independent components were extracted from the low-pass filtered MEG data (filter cutoff: 25 Hz). The obtained independent components were visually inspected and, based on their time-course and topography, those corresponding to heartbeat, eyeblink, and eye movement-related artifacts were identified. These artifact-containing components were then subtracted from the full-rank MEG signals. MEG data segments corresponding to each experimental condition were then extracted and concatenated.

#### CTS analysis

To quantify the accuracy of the cortical representation of the attended speech (i.e., CTS), we computed the coherence between the MEG signals and the speech temporal envelope of the target stream (i.e., the narrator’s voice), for each experimental condition. The speech temporal envelope was obtained as done for the generation of the vibrotactile signal (*Methods: Stimuli*). Coherence was computed based on the preprocessed MEG data split into 2 s epochs with a 1.6 s epoch overlap, which afforded a frequency resolution of 0.5 Hz^79^. Epochs exceeding 5 pT (magnetometer) or 1 pT (gradiometer) in at least one sensor were classified as artifact-containing and excluded from further analysis. Further analyses were conducted on coherence at 0.5 Hz (phrasal CTS) and 4–8 Hz (syllabic CTS).

For each condition, participant, and CTS frequency (phrasal or syllabic), a coherence map was first generated at the sensor-level using the gradiometer pair signals^80^. Then, for a more precise localization of the underlying CTS cortical sources, coherence maps were estimated at the source level. First, individual MRI volumetric segmentation was performed using the FreeSurfer software suite ^81^ (Martinos Center for Biomedical Imaging, Boston, MA, RRID:SCR_001847), after which a manual MEG-MRI co-registration was done using the digitized anatomical fiducial landmarks and the head-surface points (MRIlab, MEGIN, Finland). A non-linear transformation of the individual MRIs to the Montreal Neurological Institute (MNI) template brain was then computed using SPM8 (Wellcome Centre for Human Neuroimaging, London, UK, RRID:SCR_007037, http://www.fil.ion.ucl.ac.uk/spm/) and used to project a homogeneous 5-mm grid sampling the MNI brain onto individual MRIs. At each grid point, the currents were modeled as three orthogonally oriented dipoles. The forward model was subsequently computed using the single-layer Boundary Element Method (BEM) approach implemented in MNE-C (Martinos Centre for Biomedical Imaging, Massachusetts, USA)^82^ and reduced to its first two first principal components given the insensitivity of MEG to currents radial to the skull^17^. Lastly, the minimum-norm estimation (MNE) inverse solution^83^ was computed using a MNE regularization parameter based on the consistency condition^84^. No explicit depth bias correction was done, as coherence values are unaffected by bias correction. Coherence maps at the source level were finally created for each subject, experimental condition and CTS frequency using the dynamic imaging of coherent sources approach^85^ with the MNE inverse solution. Given that all maps were inherently coregistered to the MNI space, the group-level coherence maps for each condition and CTS frequency were obtained by averaging across participants.

#### Functional connectivity analysis

We conducted seed-based functional connectivity for the conditions and hemisphere which presented significant CTS values differences (i.e., ATc vs. A-only in multi-talker noise; right STAC). Phase synchronization was measured using the phase locking value (PLV)^38^; this connectivity measurement was chosen given its robustness to variations in signal amplitude and the demonstrated involvement of phase synchrony in large-scale (i.e., brain regions at >2 cm apart) perceptual brain integration^38,86,87^. In practice, the preprocessed MEG data for each condition was first bandpass filtered into narrow 1 Hz-wide frequency bands. A boxcar window was applied, with cutoff frequencies set to predefined limits ranging from 0.5 Hz to 40 Hz. This filtering process ensured that only the frequency ranges of interest were retained. The filter featured a transition width of 0.01 Hz at the cutoff frequencies, providing precise control over the definition of the passband. MEG noise covariance estimation was performed using a 5-minute recording of empty-room data prior to MNE-based source reconstruction. Next, brain source activity was reconstructed for each condition and within each frequency band with MNE. Seed-based functional connectivity was then measured using PLV with signal orthogonalization for spatial leakage correction^88^ from seed to 154 nodes of a dense network-based parcellation of the human brain composed obtained in a former meta-analysis of functional MRI data^89,90^. Each dipole time series was projected onto the direction of its maximum variance^91^ and the Hilbert transform was applied to derive its analytic signal. The seed was placed at the mean coordinates of the right STAC loci exhibiting significant CTS across the conditions with audible speech. The obtained between-nodes PLV values were averaged within five specific frequency bands (phrasal-delta: 0.5–1, syllabic-theta: 4–8 Hz, alpha: 8–12 Hz, low-beta: 12–21 Hz and high-beta: 21–30 Hz). Of note, each pairwise PLV was computed by using both STAC and the other node as seeds in turn and then averaged (i.e., transpose symmetrisation)^92^, to avoid possible asymmetries arising after pairwise orthogonalization.

### 7. Statistical analyses

#### Behavioral performance analysis

Participants’ speech comprehension was analyzed using a logistic mixed-effects regression analysis with the experimental condition as the fixed factor and a by-subject random intercept. This analysis was conducted in R Statistical Software^93^ (v4.1.0, https://www.R-project.org/), using the *glmer* function from the *lme4*^94^ package. Binary (yes/no) comprehension responses (correct = 1, incorrect = 0) were used as the dependent variable. The significance of fixed effects was assessed using likelihood ratio tests, with Chi-square distribution test statistics approximation. Since the model was binomial, model estimates were obtained on the log-odds scale using the *emmeans* package. For post hoc comparisons of estimated marginal means, planned one-tailed contrasts were grouped a priori into comparisons in quiet, noise and matched quiet-noise comparisons, using Wald tests for pairwise contrasts. Multiplicity correction was done using the multivariate *t* adjustment.

#### Assessment of CTS local maxima significance

A non-parametric permutation test^95^ was used to assess the statistical significance of the CTS local maxima identified in the group-averaged coherence maps, for each experimental condition and CTS frequency (phrasal and syllabic). First, subject-level and group-level *null* CTS maps were generated by computing the coherence between the original MEG signals and a time-shifted version of the attended speech signal in each video. The time-shift involved swapping the first and second halves of each recording, which would practically disrupt any genuine coupling between the attended speech and brain signals, while preserving the spectral properties of the original speech signal. The exact temporal shift applied aligned with a pause in the speech signal, ensuring continuity. Next, group-averaged difference maps were generated by subtracting *genuine* from *null* group-averaged CTS maps; under the null hypothesis, no difference between the two should exist, regardless of the experimental condition. Therefore, the labeling of *genuine* or *null* maps would be interchangeable prior to computing the difference map^95^. To reject this null hypothesis and determine the significance level of the observed difference map, the permutation-based sample distribution of the maximum absolute value of the difference maps was obtained from 1000 permutations; the use of this maximum value inherently corrects for multiple comparisons between coherence values. The statistical threshold for significance (*p* < 0.05) was then set as the 95^th^ percentile of this distribution^95^. Finally, all local maxima of CTS exceeding this threshold were interpreted as brain regions exhibiting statistically significant CTS. As permutation tests may be too conservative for voxels beyond those showing the maximum observed statistic^95^ and because CTS can present hemispheric imbalance^2,35^, we conducted the steps described above for each hemisphere separately. After identifying the coordinates of the local maxima obtained in the group-averaged CTS maps for each experimental condition, individual CTS values were extracted within a 10 mm sphere centered on the group-level coordinates.

#### Differences in CTS between the experimental conditions

Syllabic and phrasal CTS values extracted from the significant STAC local maxima were analyzed using linear mixed-effects regression analyses using the *lme4*^94^ package’s *lmer* function in the R software^93^. The experimental condition and hemisphere were set as fixed factors, and a by-subject intercept was used. Prior to the analysis, outlier fixing was done by removing CTS values deviating more than 2.5 s.d. from the distribution and setting them to the mean ± 2.5 s.d. Mixed modelling residual normality was checked with residual Q-Q plots and Kolmogorov-Smirnov testing, and model fitting was done using maximum likelihood estimation. Fixed effects were evaluated using F-tests based on Type III sums of squares. Satterthwaite’s method was used to compute the degrees of freedom for the main model’s fixed effects, while Kenward-Roger’s method was applied to the degrees of freedom for the post-hoc pairwise comparisons. Two-tailed post-hoc comparisons of the estimated marginal means obtained using the *emmeans* package were performed using paired *t*-tests with multivariate *t* multiple comparison correction.

#### Behavioral relevance of CTS vibrotactile enhancement

To determine whether the improvement in CTS values induced by the vibrotactile input was associated with changes in speech comprehension, we first computed a comprehension score. This was done by calculating the proportion of correct answers to the four content questions associated with the ATc experimental condition. For the CTS values, to better isolate the effect of supplemental non-auditory input and mitigate individual baseline auditory variability in unimodal (i.e., A-only) processing, we computed a benefit score for each hemisphere by subtracting the CTS values in the unimodal condition from those in the bimodal condition (CTS_benefit_= CTS_AT_– CTS_A_), analogously to prior investigations of audio-visual CTS enhancement^96^. Given the data count (*n* = 30) and the ordinal nature of the behavioral data (i.e., 0%, 25%, 50%, 75%, or 100%), this comprehension–CTS benefit correlation was computed using one-tailed Kendall’s tau rank correlation coefficient. All other correlational analyses were performed using one-tailed Pearson’s correlation coefficient.

#### Functional brain connectivity

To investigate between-condition differences in seed-based functional connectivity, group-averaged data was contrasted using mass-univariate unpaired *t* tests (one per node-to-node connection and frequency band) deployed in a non-parametric permutation test based on a maximum statistic approach^95^. Similarly to the CTS significance computation, this method tests the null hypothesis of no connectivity difference between conditions by permuting subject-level condition labels and creating a difference map. Here, 2000 permutations were performed to generate the distribution of the contrast coefficients of the maximum absolute values across all the connections in the difference map. The significance threshold was then established by identifying the 95^th^ percentile of the obtained null distribution. Observed *t*-statistics exceeding this threshold (calculated separately for each frequency band) were interpreted as significant functional connectivity differences between experimental conditions.

## Data and code availability

Data and code supporting the current results will be provided upon reasonable request to the corresponding author and after the approval of the institutional authorities (Hôpital Universitaire de Bruxelles and Université libre de Bruxelles).

## Author contributions

Conceptualization: I.S.R., J.B, X.D.T.; Methodology: I.S.R., M.B., V.W., J.B., X.D.T.; Software: I.S.R., M.B., V.W.; Formal analysis: I.S.R.; Investigation: I.S.R.; Resources: V.J., X.D.T.; Data curation: I.S.R.; Writing—original draft: I.S.R.; Writing—review & editing: M.B., V.W., V.J., J.B., X.D.T.; Visualization: I.S.R.; Supervision: J.B., X.D.T.; Funding acquisition: X.D.T.

## Acknowledgements

I.S.R. is supported by the Fonds pour la formation à la recherche dans l’industrie et l’agriculture (FRIA, Fonds de la Recherche Scientifique (FRS-FNRS), Brussels, Belgium). X.D.T. is Clinical Researcher at the FRS-FNRS. This research project has been supported by the Fonds Erasme (Research convention “Les Voies du Savoir”, Brussels, Belgium) and the FRS-FNRS (Research credit: J.0035.23). The MEG project at the Hôpital Universitaire de Bruxelles is financially supported by the Fonds Erasme (Research convention “Les Voies du Savoir”). The PET-MR project at the Hôpital Universitaire de Bruxelles is financially supported by the Association Vinçotte Nuclear (AVN, Brussels, Belgium). The authors would like to thank Aalto NeuroImaging (Aalto University, Espoo, Finland) for providing the vibrotactile stimulator used in the study.

